# SLRfinder: a method to detect candidate sex-linked regions with linkage disequilibrium clustering

**DOI:** 10.1101/2024.03.11.584532

**Authors:** Xueling Yi, Petri Kemppainen, Juha Merilä

## Abstract

Despite their critical roles in genetic sex determination, sex chromosomes remain unknown in many non-model organisms. In contrast to conserved sex chromosomes in mammals and birds, studies of fish, amphibians, and reptiles have found highly labile sex chromosomes with newly evolved sex-linked regions (SLRs). These labile sex chromosomes are important for understanding early sex chromosome evolution but are difficult to identify due to the lack of Y/W degeneration and SLRs limited to small genomic regions. Here we present SLRfinder, a method to identify candidate SLRs and labile sex chromosomes using linkage disequilibrium (LD) clustering, patterns of heterozygosity, and genetic divergence. SLRfinder does not rely on specific sequencing methods or reference genomes and does not require phenotypic sexes which may be unknown from population sampling, although sex information can be incorporated to provide additional inference on candidate SLRs. We tested SLRfinder using various published datasets and compared it to SATC, a method that identifies sex chromosomes based on the depth of coverage and also does not require phenotypic sex. Results show that SATC works better on conserved sex chromosomes (e.g., in African leopards), whereas SLRfinder outperforms SATC in analyzing labile sex chromosomes (e.g., in nine-spined sticklebacks and chum salmon). Since SLRfinder primarily relies on LD clusters, it is expected to be most sensitive to the SLRs harboring structural variants (e.g., inversion) due to strongly reduced recombination rates in heterozygotes. SLRfinder provides a novel and complementary approach for identifying SLRs and uncovering additional sex chromosome diversity in nature.

## Introduction

Sex chromosomes play critical roles in genetic sex determination and yet remain unknown in many non-model organisms. Early studies in mammals and birds have demonstrated highly conserved and heteromorphic (i.e., having different morphologies) sex chromosomes with conserved sex-determining genes and degenerated Y or W chromosomes. On the other hand, accumulating studies have found less conserved but much more labile sex chromosomes that may be different between closely related lineages in fish, amphibians, and reptiles (Dufresnes et al., 2015; Jeffries et al., 2018; Yi et al., 2024). These labile sex chromosomes tend to be homomorphic (i.e., sex chromosomes having indistinguishable morphologies) and are featured by little or no degeneration, low inter-sex differentiation, variable sex-determining genes, and sex-determining regions restricted to narrow genomic regions. These features make labile sex chromosomes and their sex-determining regions difficult to identify using traditional methods such as karyotyping and PCR of conserved sex-determining genes (Palmer et al., 2019; Tree of Sex Consortium, 2014). However, labile sex chromosomes likely represent early evolutionary stages of sex chromosome evolution and their study is critical for our understanding of sex chromosome evolution (Blaser et al., 2014; Furman et al., 2020; Perrin, 2021; Vicoso, 2019). Therefore, additional work is needed to identify labile sex chromosomes and their sex-determining regions in non-model species.

Recently, several methods have been developed to help identify sex chromosomes and their sex-determining regions in non-model species, but these methods mostly work for conserved sex chromosomes and are limited to certain types of sequencing data. For example, RADSex (Feron et al., 2021) was developed to identify sex determination systems (i.e., XX/XY or ZZ/ZW) and sex-linked markers of labile sex chromosomes specifically from restriction site-associated DNA sequencing (RADseq) data, and Pooled Sequencing Analysis for Sex Signal (PSASS ver. 3.1.0; https://github.com/SexGenomicsToolkit/PSASS) was developed to detect sex-linked signals using pooled sequencing data from males and females (e.g., in Kitano et al., 2023). These methods are not applicable to whole-genome sequencing (WGS) data which has been increasingly used in studies of non-model populations. In addition, these methods require known phenotypic sexes which may not be available in non-invasive sampling or may be difficult to identify in individuals that are not sexually mature or have limited or no sexual dimorphism. FindZX (Sigeman et al., 2022) was developed to detect sex chromosomes using WGS data. This method has been applied to diverse systems including both conserved and labile sex chromosomes, and it can work on very small sample sizes (Sigeman et al., 2022). However, this method also relies on known phenotypic sexes, and it requires a reference genome of the homogametic sex (i.e., XX female or ZZ male) which may not be available or may be unknown if the sex determination system is unclear. SATC (Sex Assignment Through Coverage; Nursyifa et al., 2022) was developed to jointly identify sex chromosomes and genetic sex using WGS data. This method does not require known phenotypic sexes, but it assumes that only X/Z scaffolds are assembled in the reference genome, which is practically the same as requiring a reference genome of the homogametic sex. In addition, the available methods are mostly based on sequencing depth (RADSex and SATC) or depth and heterozygosity (PSASS, FindZX), but many studies have shown that depth may not differ between sexes on labile sex chromosomes which are homomorphic with narrow sex-determining regions (Jeffries et al., 2022; Yi et al., 2024). Therefore, new methods are needed to help identify labile sex chromosomes in non-model populations.

A previous study has shown that LD can be a signal for detecting sex-determining regions (McKinney et al., 2020). Here we present a method (herein SLRfinder) to identify candidate sex-linked regions (SLRs) among clusters of highly correlated SNPs (LD clusters) based on the differentiation in heterozygosity and the genetic differentiation captured by Principal Component Analysis (PCA). The identified candidate SLRs should include the sex-determining region and its linked genomic regions on sex chromosomes, as well as possible rare sex-linked autosomal regions. However, the sex-determining region is expected to have the strongest signal of LD due to recombination suppression between sex chromosomes, and the clearest genetic divergence between sexes captured by PCA. The heterozygosity difference in the sex-determing region can be used to indicate homogametic and heterogametic sexes. SLRfinder is expected to outperform the coverage-based methods in identifying young sex chromosomes, and it does not rely on specific types of sequencing methods or reference genomes. Like SATC, SLRfinder does not require phenotypic sexes, although known sex information can be incorporated as a complementary filtering to provide additional inference for candidate sex chromosomes.

Below we describe the workflow of SLRfinder and its application to published datasets of various taxa having identified labile sex chromosomes, including nine-spined sticklebacks (*Pungitius pungitius*), chum salmon (*Oncorhynchus keta*), and guppies (*Poecilia reticulata*). We also tested the effectiveness of SLRfinder in conserved sex chromosomes using a dataset of African leopards (*Panthera pardus*). In addition, we compared the performance of SLRfinder to SATC, the only other method that does not require known phenotypic sexes. Results show that, as expected, SATC only worked on conserved sex chromosomes and might yield wrong sex inferences when using a reference genome of the heterogametic sex. On the other hand, SLRfinder does not rely on specific types of reference genomes and it outperforms SATC in analysing labile sex chromosomes, especially when the SLR is associated with genomic inversions. Since SLRfinder and SATC are based on independent signals (i.e., LD and heterozygosity versus depths of coverage), they are complementary to each other and should thus be considered jointly to maximize the ability to identify sex chromosomes in non-model species.

## Materials and Methods

### Identify LD clusters from VCF inputs

The workflow of SLRfinder is summarized in **Figure 1**. The input data is a VCF file of filtered biallelic single nucleotide polymorphisms (SNPs) from populations genotyped using WGS or reduced-representation sequencing methods (e.g., RADseq). LD is estimated in VCFtools (Danecek et al., 2011) as squared coefficient of correlation (*r^2^*) between pairs of loci within windows of 100 SNPs (--geno-r2 --ld-window 100). In a network analytical framework (Kemppainen et al., 2015), LD clusters are identified by only considering LD values (edges) with *r^2^*>*min_LD* (default 0.85) between pairs of loci (nodes). The resulting edge list (each row corresponding to a locus pair with *r^2^*>*min_LD*) is used to generate a “graph object” using the function *graph.edgelist* from the package igraph (Csardi et al., 2006) in R v4 (R Core Team, 2022). The graph object is further decomposed into separate LD clusters that are not connected by any edges using the function *decompose.graph*. The resulting LD clusters are defined by their weakest edges such that only loci belonging to the same cluster can be connected by *r^2^*>*min_LD* (i.e. single linkage clustering). Finally, only clusters with a minimum of *min.cl.size* loci (default 20) are retained for downstream analyses.

**Figure 1.**
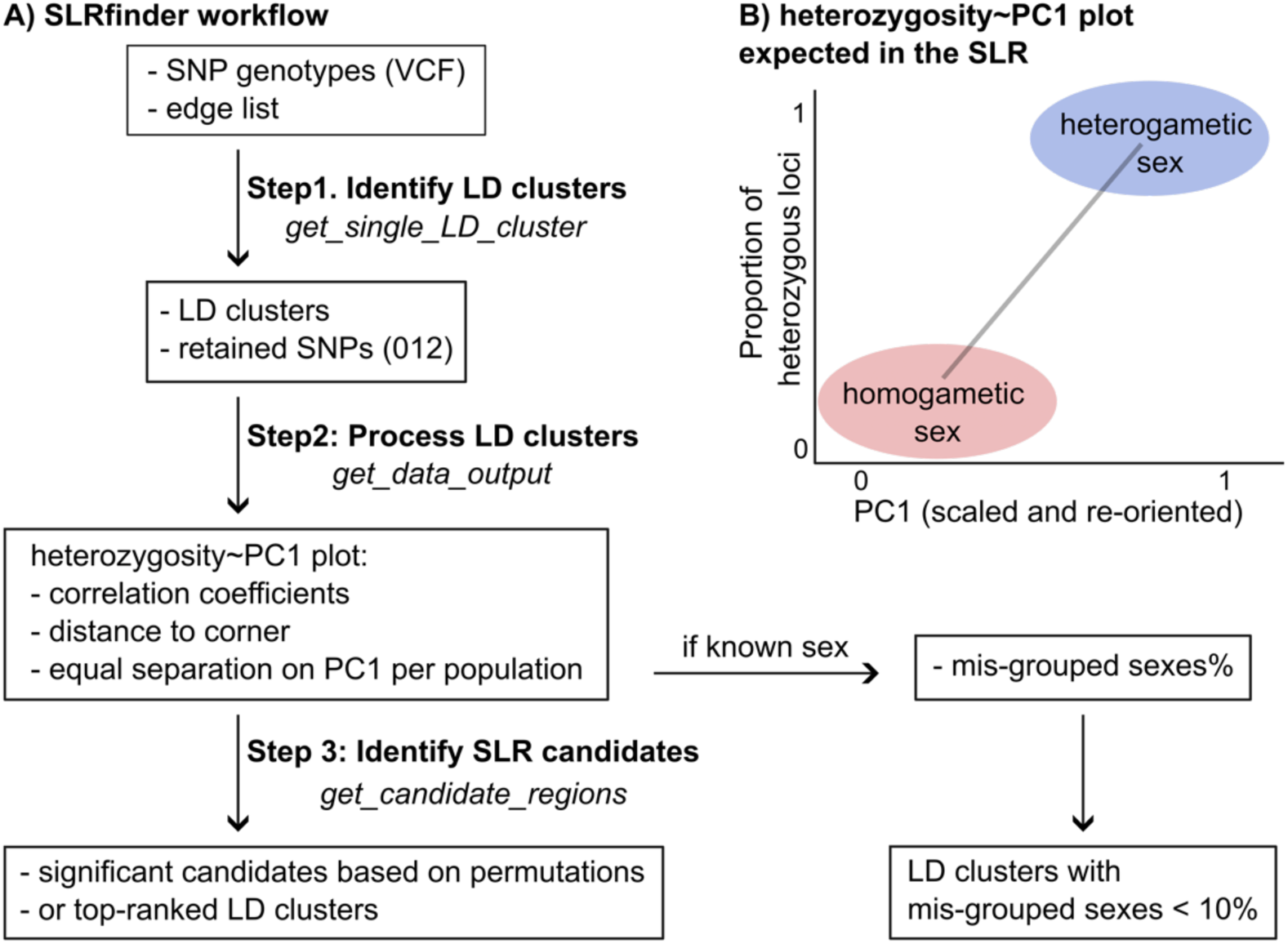
The workflow of SLRfinder. **A)** The three major steps of SLRfinder. **B)** Illustration of the expected heterozygosity~PC1 plot in the SLR.

### Estimating heterozygosity and conducting PCA in each LD cluster

All SNPs from each identified LD cluster are used to conduct a PCA using the R package SNPRelate (Zheng et al., 2012) and estimate the observed heterozygosity as the proportion of heterozygous SNPs (coded as 1 in the 012 file) in the non-missing SNPs genotyped in each individual. A linear model is fitted to regress the estimated individual heterozygosity on scaled PC1 (also polarized if the original relationship is negative). The heterozygosity~PC1 plots are expected to show no grouping pattern in most LD clusters, three groups in a triangular shape representing three genotypes in autosomal inversions (Ma & Amos, 2012), and two groups corresponding to the homogametic sex (bottom-left corner) and the heterogametic sex (top-right corner) in SLRs (**Fig. 1B**). Accordingly, candidate SLRs are expected to have stronger association between heterozygosity and PC1, and stronger inter-sex genetic divergence than population structure captured on PC1. Based on these expectations, we estimate the adjusted R-squared values of the linear regressions and the ꭓ^2^ goodness-of-fit tests on an equal separation of samples in each population on PC1 under the assumption of an equal sex ratio and random sampling. Smaller ꭓ^2^ statistics indicate that all populations include individuals from both groups, therefore indicating potentially stronger inter-sex differentiation than population structure in this region. If in some cases skewed sex ratios are expected, the expected probabilities of sampling the heterogametic sex and the homogametic sex can be provided to get more accurate ꭓ^2^ estimates. In addition, we estimated the scaled Euclidean distance between each individual and its nearest corner individual (i.e., the individuals having the highest or lowest heterozygosity, and if equal heterozygosity the highest or lowest scaled PC1 scores). Assuming no recombination, candidate SLRs are expected to have a clear and strong separation between groups on the heterozygosity~PC1 plot, resulting in shorter Euclidean distances with smaller variance.

### Identify SLR candidates

Candidate SLRs are identified among LD clusters based on their ranks of the estimated parameters. A LD cluster is ranked higher (i.e., more likely to be a SLR) if it has more SNPs, stronger heterozygosity~PC1 regression, smaller variation of the Euclidean distance (i.e., better grouping on the heterozygosity~PC1 plot), and smaller ꭓ^2^ statistic (i.e., roughly equal separation of individuals per population on PC1). The summed ranks of these parameters are permuted (default 10000 times) among LD clusters to generate a null-distribution of the summed ranks and estimate how often the permuted values are lower than the observed value (i.e., the *p*-value) of each LD cluster. In addition, we correct potential *p*-value inflation using genomic control (Devlin et al., 2001; Devlin & Roeder, 1999). Briefly, the −log10(*p*) values are divided by the inflation factor (*λ*) estimated as the linear slope in a quantile-quantile plot between the observed −log10(*p*) and those expected under the null-hypothesis of a uniform distribution of *p-*values. Significant (adjusted *p*-value < 0.05) candidates (or the five top-ranked LD clusters if no significance) are reported with their heterozygosity~PC1 plots.

Although SLRfinder does not require phenotypic sexes, known sex information can be incorporated to filter LD clusters where the two sexes are fully separated on the heterozygosity~PC1 plot, which can provide additional inference on candidate sex chromosomes. To do this, we estimate the percentage of sexed individuals that are likely placed in the wrong group (i.e., the minority sex in a group is regarded as misplaced), and filter the LD clusters that have less than 10% mis-placed individuals (allowing for rare phenotypic misidentifications).

### Test of SLRfinder using published datasets

To test the efficiency and accuracy of SLRfinder, we applied it to published empirical datasets of various species (**Table S1**). First, we applied SLRfinder to the nine-spined stickleback (*Pungitius pungitius*) lineages that have different sex chromosomes. Previous studies have shown that the non-European and Eastern European (EL) lineages have heteromorphic sex chromosomes identified as LG12, whereas the Western European lineage (WL) have homomorphic sex chromosomes identified as LG3, and two UK populations have unidentified sex chromosomes (Dixon et al., 2019; Natri et al., 2019; Yi et al., 2024). The WGS data of nine-spined sticklebacks were published in a previous study (Feng et al., 2022) and available on ENA (project PRJEB39599). The raw sequencing data were re-mapped to the version 7 reference genome of *Pungitius pungitius* (GCA_902500615.3; Kivikoski et al., 2021) using bwa-mem in BWA v0.7.17 (Li, 2013), sorted and indexed using SAMtools version 1.16.1 (Danecek et al., 2021), and genotyped by Genome Analysis Toolkit (GATK) following the best practice protocol (Depristo et al., 2011; Van der Auwera et al., 2013). Biallelic SNPs were extracted using the commands -m2 -M2 -v snps –min-ac=1 in BCFtools (Li, 2011) and data mapped to unassembled contigs were removed. The SNP genotypes were split into four datasets representing the WL, the EL, the non-European lineage, and the UK lineage. Each dataset was further filtered in VCFtools by quality (--minGQ 20 --minQ 30), missing data (--max-missing 0.75), and minor allele frequency (--maf 0.15) before analysed by SLRfinder. The same filtering was used below in the other test datasets. Phenotypic sexes are known in one EL and one WL population and were provided to SLRfinder.

Next, we applied SLRfinder to chum salmon (*Oncorhynchus keta*) whose sex chromosomes have been identified as LG15 in studies using RADseq (McKinney et al., 2020) or WGS data (Rondeau et al., 2023). We re-analysed both datasets using SLRfinder. The WGS data were mapped to the newly assembled male reference genome of *Oncorhynchus keta* (GCF_023373465.1) and the VCF file of genotyped bi-allelic SNPs were downloaded from the corresponding publication (Rondeau et al., 2023) and filtered before being analysed by SLRfinder. In addition, to test the potential influence of different reference genomes, we downloaded the raw WGS data from NCBI (BioProject PRJNA556729), mapped them to a female reference genome (GCF_012931545.1), and genotyped and filtered SNPs in the same way described above. To test the application of SLRfinder on reduced-representation sequencing data, we also re-analysed the RADseq data published in (McKinney et al., 2020). The demultiplexed raw sequencing data were downloaded from NCBI (BioProject PRJNA611968) and mapped to the male reference genome (GCF_023373465.1) using bwa-mem. The mapped reads were sorted, indexed, and marked with duplicates using SAMtools and genotyped using the program *ref_map.pl* with default settings in Stacks 2.65 (Rochette et al., 2019). The genotyped data were further filtered using the program *populations* by minor allele frequency (--min-maf 0.15) and missing data (-R 0.75), and the ordered genotypes were output in the VCF format. We did not output a single SNP per stack locus as the following analyses are based on the information of linkage disequilibrium. The output VCF file was analysed by SLRfinder using a lower threshold for detecting LD clusters (*min_LD*=0.2, *min.cl.size*=5) due to lower SNP density in RADseq data. Phenotypic sexes are known for the WGS dataset (Rondeau et al., 2023) but not the RADseq dataset (McKinney et al., 2020).

We also applied SLRfinder to datasets of guppies (*Poecilia reticulata*) whose sex chromosomes have been identified as the LG12 with two SLR candidates indicated using a newly assembled male reference genome (Fraser et al., 2020). The raw WGS data of previously studied populations (Fraser et al., 2020; Kü Nstner et al., 2016) were downloaded from NCBI (BioProject PRJEB10680, PRJNA238429) and mapped separately to the male reference genome (GCA_904066995.1) and a female reference genome (GCA_000633615.2) to test potential impacts of using different references. Data mapping, genotyping, and SNP filtering were done in the same way as in nine-spined sticklebacks. Phenotypic sexes are known for these individuals (Fraser et al., 2020) and were provided to SLRfinder.

Lastly, we applied SLRfinder to African leopards (*Panthera pardus*) which have conserved sex chromosomes. Due to computational constraints, we only analysed the WGS data of 26 individuals published in a previous study (Pečnerová et al., 2021). The raw data were downloaded from NCBI (BioProject PRJEB41230) and mapped to a scaffold-level female reference genome of *Panthera pardus* (GCF_001857705.1). Data mapping, genotyping, and SNP filtering were done in the same way as in nine-spined sticklebacks. Because sample populations are not provided for the raw sequencing data on NCBI, we assigned these individuals into genetic populations based on PCA using separately filtered biallelic SNPs (--minGQ 20 --minQ 30 --maf 0.05 --max-missing 0.8). No phenotypic sexes were provided and the genetic sexes inferred by SATC (see below) were used as the sex information in SLRfinder.

### Identifying sex and SLRs using SATC

We also compared the effectiveness of SLRfinder to SATC (Nursyifa et al., 2022) using the above datasets, excluding the salmon WGS data mapped to the male reference genome because this dataset was a VCF file downloaded from the previous publication (Rondeau et al., 2023) and the bam files were not available. To run SATC, the depth of coverage was calculated by SAMtools-idxstats using the mapped and duplicates-marked individual bam files. Then the idx files were processed by SATC with default settings which filter scaffolds by minimum 100kb, normalize length by the five longest scaffolds, and identify sex scaffolds by the Gaussian model.

### Testing the power of SLRfinder using different sample sizes and sex ratios

To assess the statistical power of SLRfinder, we applied it to subsets of the WL and EL nine-spined stickleback datasets where we varied the number of individuals or populations and tested uneven sex ratios. To test effects of sample sizes, we first randomly selected 3-5 individuals per population while including all populations in the WL or EL dataset. Then we randomly selected 1-5 WL or EL populations while including all individuals from the selected populations. To test effects of sex ratios, we used the previously identified genetic sexes of these individuals (Yi et al., 2024) and only included the seven WL populations and the 24 EL populations that have at least 4 individuals per sex. We randomly selected 2 individuals per sex per population (even sex ratio), or 1 individual from one sex and 3 from the other in each population (sex ratios 1:3 or 3:1). To test sex ratios 1:2 or 2:1, we randomly selected 9 individuals from one sex and 19 from the other across WL populations, and 32 individuals from one sex and 64 from the other across EL populations. To test sex ratios 1:10 or 10:1, we randomly selected 3 individuals from one sex and 25 from the other across WL populations, and 9 individuals from one sex and 87 from the other across EL populations. We also tested extreme scenarios where only one sex was sampled in the dataset. The subset VCF files of the selected individuals were filtered and processed by SLRfinder as described above. We first used the default expectation of an equal sex ratio in all tests. When the true SLR was not included in top-ranked candidates, we further modified the parameters to rank and the expected sex ratios in the ꭓ^2^ tests to see if SLRfinder results can be improved.

## Results

### SLRfinder analyses of nine-spined sticklebacks and chum salmon

SLRfinder successfully identified the sex chromosomes and SLRs of nine-spined sticklebacks (Table 1; Fig. 2). In the WL dataset, SLRfinder identified a single significant candidate on LG3 that highly overlaps with the previously described WL SLR (LG3:17260000-17340000 bp; Yi et al., 2024). Similarly, in the EL and non-European datasets, SLRfinder identified a single significant candidate on LG12 that highly overlaps with the previously reported EL SLR (LG12:1-16900000 bp (Kivikoski et al., 2021). The SLRfinder-inferred genetic sexes are also consistent with known phenotypic sexes and the previous identifications of genetic sex (Yi et al., 2024). The UK dataset did not generate significant candidates, possibly due to the limited power of SLRfinder on small sample sizes (see below). However, none of the top-ranked candidates were located on LG12 or LG3 (Table S1), consistent with the previous findings that sex chromosomes of the UK populations are likely neither LG12 nor LG3 (Yi et al., 2024). Instead, the LD clusters having the lowest adjusted *p-*values (*p*=0.2179) included a 225-bp region on LG7 and a 203-bp region on LG16 (Table 1, Fig. S1A). Additional sampling of individuals with known sexes is required to validate if these regions can separate the two sexes and to identify the yet unknown sex chromosomes of the UK populations.

**Figure 2.**
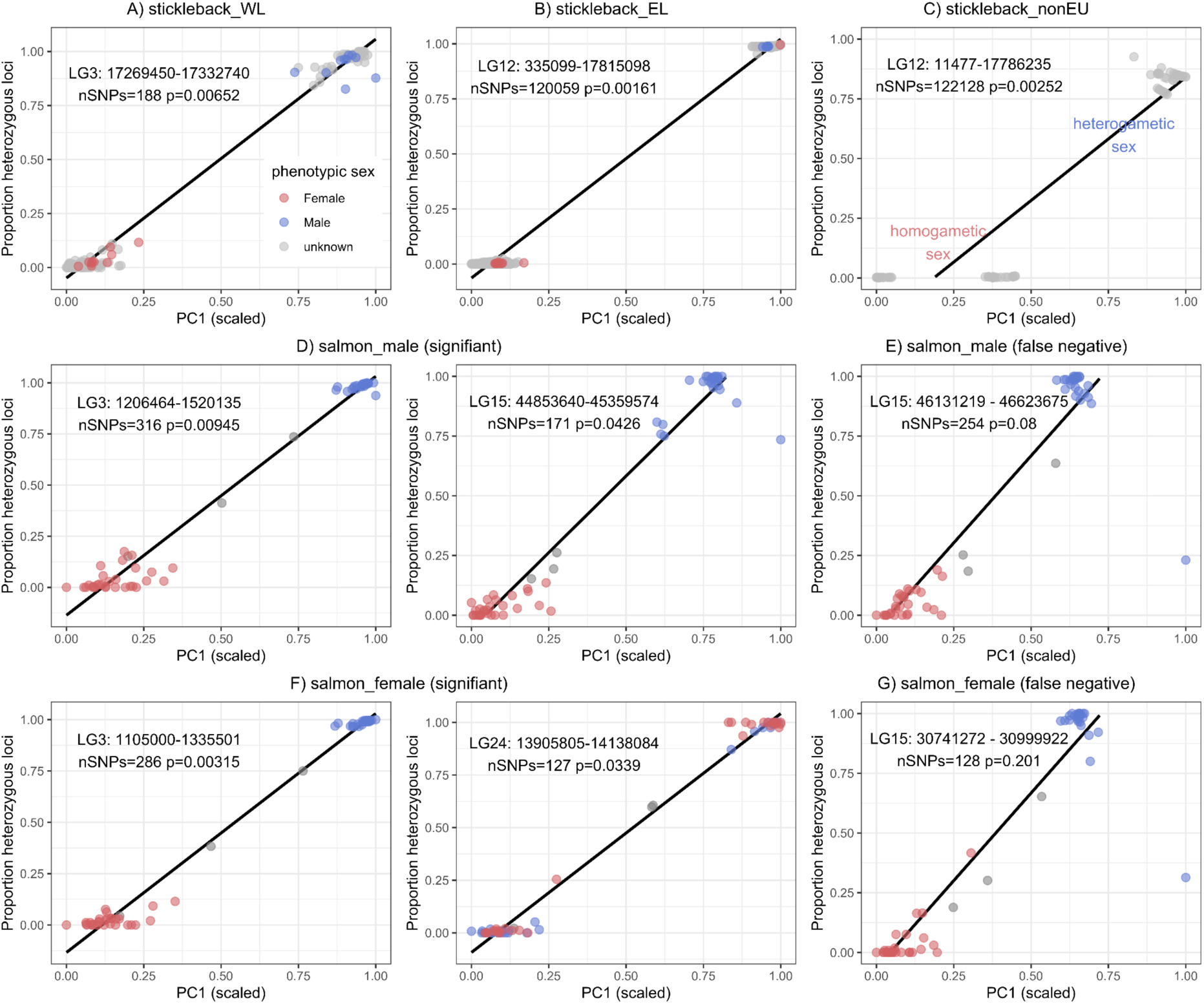
The heterozygosity-PC1 plot of the SLRfinder-identified candidates using datasets of nine-spined sticklebacks (A-C) and chum salmon (D-G). Dots represent individuals colored by the phenotypic sex. The black line represents the fitted linear regression. **B)** The single phenotypic female in the top-right group is the individual 16-f that was also found to be a genetic male in previous studies (Feng et al., 2022; Yi et al., 2024). **E & G)** The false negative clusters on LG15 detected by the sex percent filtering. Two more false negatives were detected in the salmon_female dataset but had much fewer SNPs (nSNPs of 26 and 96) and thus were not plotted.

**Table 1.** Summary of the SLRfinder results using test datasets. Sex-filtered results are the LD clusters having less than 10% misplaced sexed individuals. Ranked candidates are the LD clusters tested significant (adjusted *p* < 0.05) or, if non-significant, the clusters having the lowest adjusted *p*-value (only those with *p* < 0.5 are listed). Clusters on the known sex chromosomes are indicated in bold.

| Dataset | # Ind | # Pop | # LD cluster | Sex_filter | Rank_candidates |
| --- | --- | --- | --- | --- | --- |
| stickleback_WL | 162 | 8 | 2737 | **LG3 (1 cluster)** | **LG3: 17269450-17332740** |
| stickleback_EL | 598 | 29 | 1149 | **LG12 (6 clusters)** | **LG12: 335099-17815098** |
| stickleback_nonEU | 78 | 5 | 1329 | Sex unknown | **LG12:11477-17786235** |
| stickleback_UK | 29 | 2 | 5331 | Sex unknown | LG7: 3628961-3664806,<br>LG16: 12008573-12103926<br>($p = 0.2179$) |
| salmon_male | 59 | 11 | 25646 | LG3 (2 clusters),<br>**LG15 (2 clusters)** | LG3: 1206464-1520135,<br>**LG15: 44853640-45359574** |
| salmon_female | 59 | 11 | 28294 | LG3 (2 clusters),<br>**LG15 (3 clusters)**,<br>LG26 (2 clusters),<br>LG32 (1 cluster) | LG3:1105000-1335501,<br>LG24:13905805-14138084 |
| salmon_RAD | 288 | 6 | 1498 | Sex unknown | LG14: 53640831-53640941,<br>**LG15: 22646022-46527777**<br>($p = 0.06$) |
| guppy_female | 170 | 10 | 103 | No cluster retained. | All clusters had $p > 0.5$ |
| guppy_male | 170 | 10 | 78 | No cluster retained. | All clusters had $p > 0.5$ |
| leopard | 26 | 3 | 90 | NW_017619865.1<br>NW_017619916.1<br>NW_017619950.1<br>NW_017619951.1<br>NW_017619964.1<br>NW_017620089.1 | All clusters had $p > 0.5$ |

SLRfinder also identified the sex chromosomes and SLRs of chum salmon (Table 1; Fig. 2). When using the WGS data mapped to the male reference, SLRfinder identified LG15 and LG3 as significant candidates, both highly overlapping with the previously reported sex-associated regions (LG3:750001-1950001, LG15:40010001-46610001, and LG26:1-280001) in Genome-Wide Association Studies (GWAS; Rondeau et al., 2023). Because LG15 was inferred as sex chromosomes by independent studies using different datasets and analyses (McKinney et al., 2020; Rondeau et al., 2023), the LG3 cluster most likely represents a true sex-linked autosomal region. Interestingly, despite the complete separation between two sexes on the heterozygosity~PC1 plot of this LG3 cluster, the few individuals of unknown sex were not grouped with either sex in the LG3 cluster but were clearly grouped with females in the significant LG15 cluster which is the true SLR (Fig. 2D). Another LG15 cluster located within the previously identified SLR (LG15:40010001-46610001; Rondeau et al., 2023) was also detected by filtering the percentage of misplaced sexes. Therefore, this cluster was a false negative (*p*=0.08) with a marginal rank probably due to an outlier male individual on the heterozygosity~PC1 plot (Fig. 2E). Similarly, when using the WGS data mapped to the female reference, the autosomal LG3 cluster was identified significant and three LG15 clusters were detected by filtering the misplaced sexes but were ranked as false negatives (*p*>0.2) probably due to an outlier male that had relatively low heterozygosity (Fig. 2FG, Table 1, Table S1). On the other hand, SLRfinder identified a false positive (*p*=0.03) LG24 cluster which showed a similar pattern but did not separate two sexes on the heterozygosity~PC1 plot (Fig. 2F). When using the RADseq data, no significant candidate was identified but the true LG15 SLR was the top-ranked cluster having 319 SNPs and a marginal *p*-value of 0.06 (Table 1). This false negative result was possibly due to the sparse RADseq SNPs and loose LD filtering (*min_LD*=0.2) of this dataset which generated a weak grouping on the heterozygosity~PC1 plot (Fig. S1B).

### SLRfinder analyses of guppies and leopard

SLRfinder did not identify significant candidates using the datasets of guppies (Table 1). Previous studies have identified LG12 as the sex chromosomes of guppies (Fraser et al., 2020). However, none of the top-ranked clusters were located on LG12 (Table S1, Fig. 3A, Fig. S2A). In fact, despite a relatively large sample size (170 individuals, 10 populations), the guppy datasets were identified with very few LD clusters (Table 1) including only two LG12 clusters using the female reference genome (Fig. S2BC) and one LG12 cluster using the male reference genome (Fig. 3B), none of which showed a separation between sexes. To further investigate the signal of SLRs in guppies, we extracted SNPs located in the previously reported candidate SLRs (LG12: 4800000-5200000 bp, LG12: 24500000-25400000 bp) using the filtered VCF mapped to the male reference genome that was used to identify these SLRs (Fraser et al., 2020). For each SLR, all SNPs were used to generate the heterozygosity~PC1 plot, which showed similar heterozygosity in males and females and stronger population structure than sex differentiation on PC1 (Fig. 3CD). Therefore, these results indicate that the guppy datasets do not have the expected signal for SLRs (i.e., inter-sex differentiation in heterozygosity and stronger differentiation between sexes than population structure on PC1), which explains why SLRfinder was not able to identify these SLRs.

**Figure 3.**
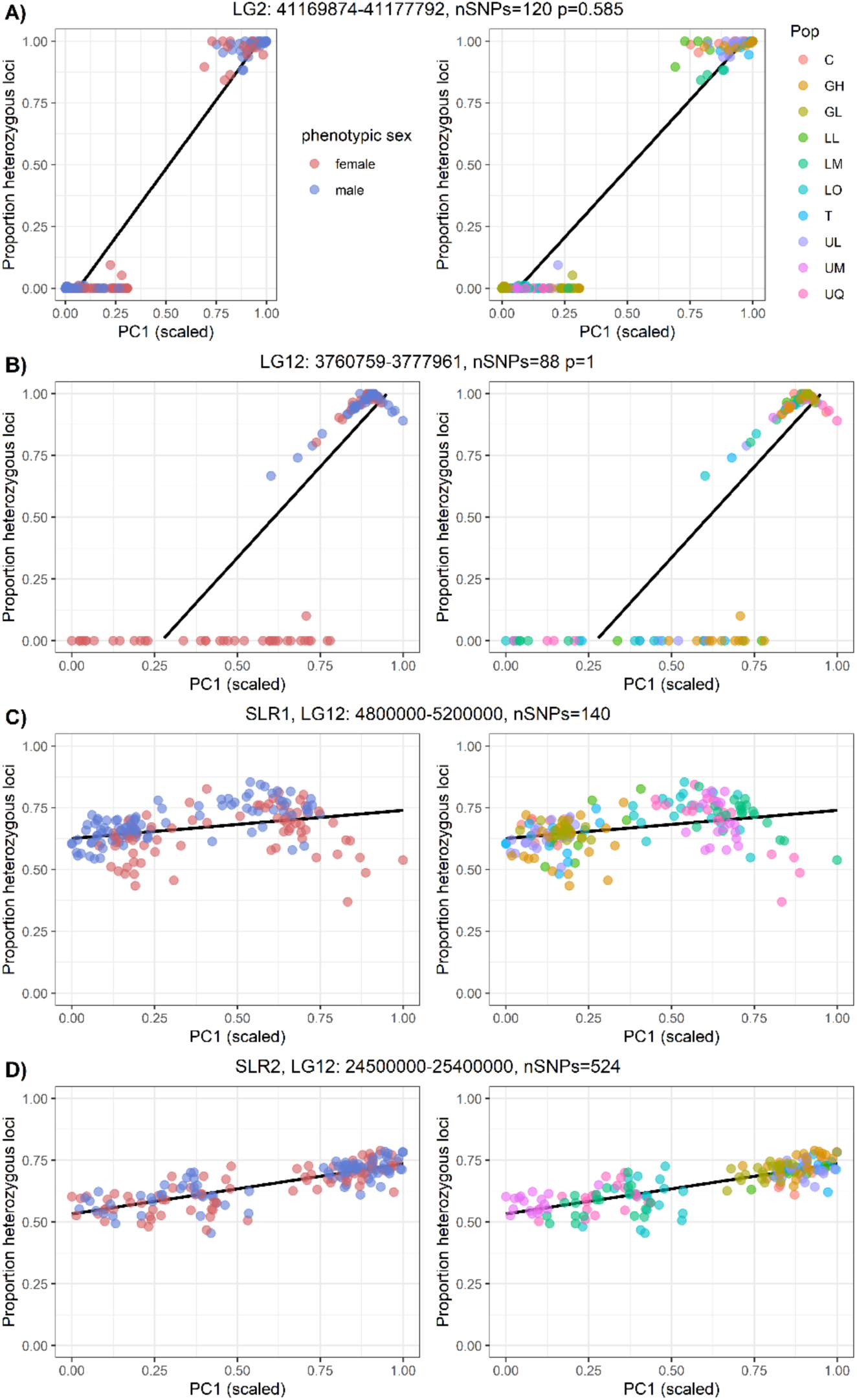
The heterozygosity~PC1 plots of the guppy dataset mapped to the male reference genome. Each dot is one individual colored by phenotypic sex (left) or population (right). **A)** The top candidate identified by SLRfinder. **B)** The single LD cluster identified on the sex chromosomes LG12. **C & D)** Plots using SNPs from the two previously reported SLR candidates (Fraser et al., 2020). The two sexes did not differ in heterozygosity, and the PC1 divergence mostly reflects population structure.

SLRfinder also did not find significant candidates in the dataset of African leopards, using the SATC-inferred genetic sex and the PCA-inferred genetic populations (Fig. S3A). However, six LD clusters were detected by filtering the misplaced sexes (Fig. S3B) and two of them were located on the scaffolds that were also identified with abnormal depth ratios in SATC (see below), indicating that these clusters are likely truly sex-linked.

### SATC analyses of test datasets

SATC could not analyze the datasets of WL sticklebacks, UK sticklebacks, chum salmon mapped to the female reference, or guppies mapped to the male reference genome. Specifically, when using the *sexDetermine* command to identify sex and sex scaffolds in these datasets, SATC reported an error of no good candidates found based on the depth of coverage, which is consistent with the previously shown lack of differentiation of depth between sexes in most of these populations (Fraser et al., 2020; Yi et al., 2024). Although SATC was able to process the chum salmon RAD data mapped to the male reference genome and the guppy dataset mapped to the female reference genome, the inferred genetic sexes were wrong and the known sex chromosomes could not be identified (Fig. 4CD), likely because the chromosome-level depth difference was small and SATC could not break down long chromosomes into small regions of SLRs.

When analyzing EL nine-spined sticklebacks, SATC inferred genetic sexes that were opposite to phenotypic sexes or the previously identified genetic sexes (Yi et al., 2024), and wrongly identified a putatively Y-linked unassembled contig (ctg7180000006428; Kivikoski et al., 2021) as X/Z-linked (Fig. 4A). This is because SATC assumes only X/Z-linked contigs in the reference genome and therefore always identifies the individuals having a higher depth of coverage on the sex-linked contigs as the homogametic sex (Nursyifa et al., 2022). When the unassembled Y-contigs are included in the reference genome, such as in the case of EL nine-spined sticklebacks (Kivikoski et al., 2021), these contigs would show strong signals of low depth in XX females while high depth in XY males, which would be misinterpreted by SATC assuming only X/Z-linked contigs. When analyzing non-European sticklebacks, SATC did not detect X/Z-linked regions and only indicated several regions with abnormal depth ratios (Fig. 4B). Interestingly, the SATC-inferred genetic sexes were consistent with results from SLRfinder (Fig. 2C) and the previous study (Yi et al., 2024) in non-European sticklebacks, except for a Canadian population (CAN-FLO) whose individuals were indicated as genetic males in SLRfinder and in the previous study but genetic females in SATC. Additional sampling with known phenotypic sexes is required to validate the sex identification of these non-European populations. Overall, these results showed limited application of SATC to the identification of labile sex chromosomes.

**Figure 4.**
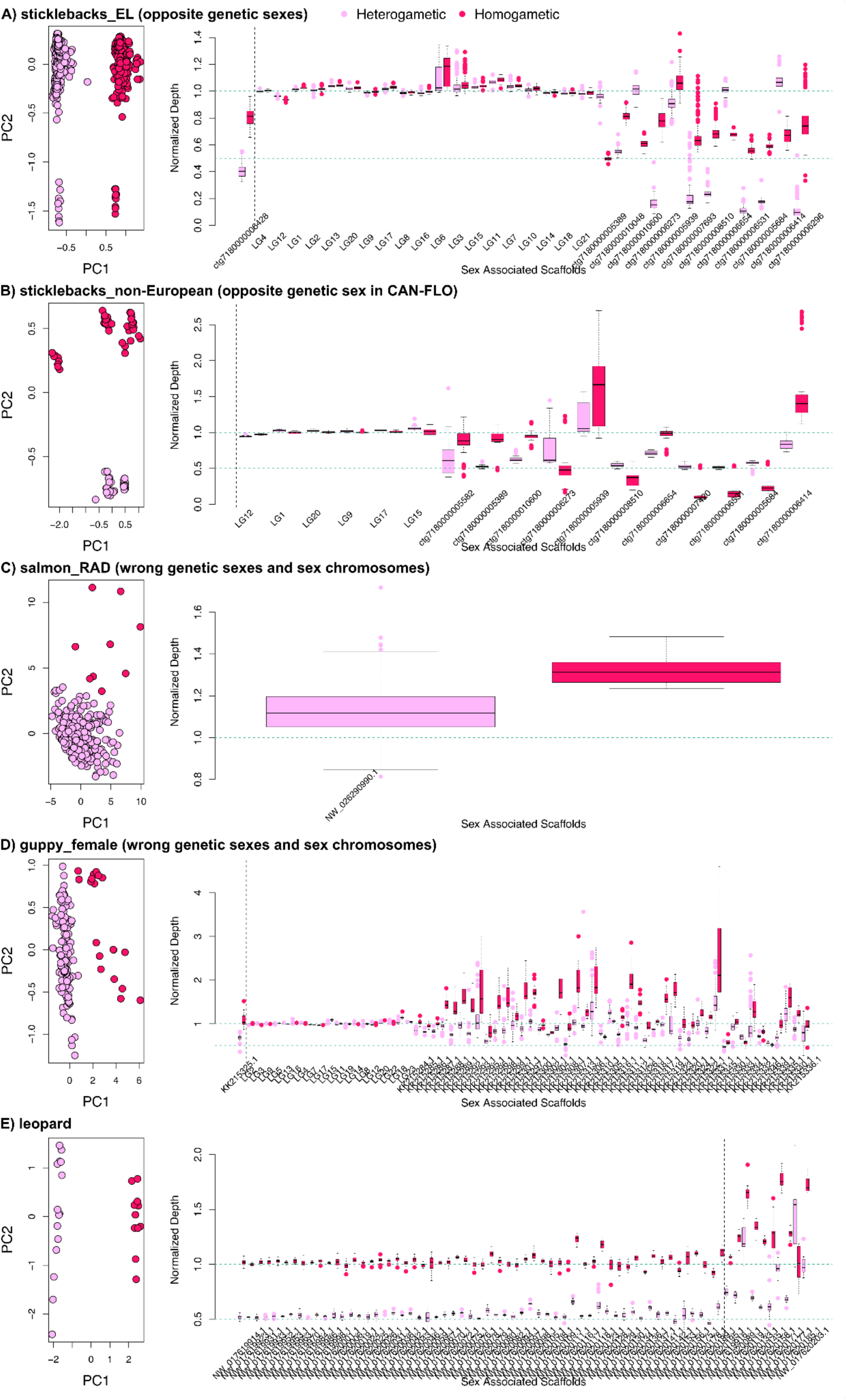
SATC results of test datasets. Colours represent the SATC-inferred homogametic (dark red) and heterogametic (pink red) sexes. The left column is the PCA plot of the normalized depth of coverage across all scaffolds and samples. The right column is the boxplot of the normalized depth of coverage of each sex from the identified sex-linked scaffolds. The vertical dashed line separates the scaffolds that passed both the t-test and ratio-based threshold (i.e., X/Z-linked, left to the line) from those that only pass the t-test (i.e., abnormal depth ratios, right to the line).

On the other hand, SATC was successfully applied to the dataset of African leopards which have conserved sex chromosomes and were mapped to a scaffold-level female reference genome. Using only 29 individuals, we identified 58 scaffolds as X/Z-linked and 8 scaffolds having abnormal depth ratios (Fig. 4E), including all of the reported sex-linked scaffolds in previous studies using the same dataset (Nursyifa et al., 2022; Pečnerová et al., 2021).

### Power tests of SLRfinder

Results of the power tests using the WL and EL sticklebacks are summarized in Table S2 and Table S3, respectively. In the WL sticklebacks, SLRfinder accurately detected the LG3 SLR as the only significant candidate when using all eight populations with three to five randomly selected individuals per population (minimum 24 individuals in total). The LG3 SLR was always identified with the lowest *p*-value when using one to five populations in the dataset, although only the test using five populations showed significance. SLRfinder identified the LG3 SLR as the significant candidate when testing the sex ratios (male:female) of 1:1 or 3:1 and with the lowest *p*-value when testing the sex ratios of 1:2, 2:1, or 10:1. No significance was found and the LG3 SLR was not identified among top-ranked candidates when testing the sex ratios 1:3 and 1:10 with the null expectation of even sex ratios. We then set the expected sex ratio as 1:3 and 1:10 and re-ran SLRfinder on these datasets, respectively. Tests of the sex ratio 1:3 with the corrected expectation and excluding the rank of number of SNPs showed that the LG3 SLR had the lowest *p-*value (Table S2), but this result was not significant (*p*=0.2) probably because few SNPs from the SLR were genotyped when few individuals of the heterogametic sex were included in the samples. However, even if using the correct sex ratio and no rank of number of SNPs, the LG3 SLR was not included in the top-ranked candidates when the sex ratio was extremely skewed (1:10 or 10:1). The LG3 SLR was detected by filtering on the percentage of misplaced sexes (based on the previously identified genetic sexes) in most of these datasets. When one sex was completely missing, neither sex filtering nor the candidate ranking could work and no false positives were found.

When applied to the EL sticklebacks, SLRfinder accurately detected the LG12 SLR as the only significant candidate when using all 29 populations with three to five randomly selected individuals per population, and when using three to five randomly selected populations (Table S3). When only one population was included, the LG12 SLR was identified with a marginal *p*-value (0.08) as the top-ranked candidate. However, when using two populations, the LG12 SLR was not included in the top-ranked candidates, indicating some uncertainty when the sample size is small and the sex ratio is uneven (around 1:3 in this case; using this sex ratio as the expectation allowed SLRfinder to add a LG12 cluster to the top-ranked candidates). When testing different sex ratios, SLRfinder identified the LG12 SLR as the significant candidate(s) using sex ratios of 1:1, 1:2, 2:1, 1:3, and 3:1. No significance was found and the LG12 SLR was not identified among the top-ranked candidates when using the most skewed sex ratios (1:10, 10:1). However, when providing the correct sex ratio of 10:1, three significant candidates were detected, including the LG12 SLR and two false positives (*p*=0.0491, Table S3). The LG12 SLR was detected by filtering based on the percentage of misplaced sexes in all datasets except for those having the most skewed sex ratios (1:10, 10:1). Again, neither sex filtering nor the candidate ranking could work when one sex was completely missing.

## Discussion

Linkage disequilibrium (LD) has been shown to be highly informative about chromosomal evolution, adaptation, and population structure (Kemppainen et al., 2015; Faria et al., 2019; Fang et al., 2020; Fang et al., 2021; Guzmán et al., 2021), and has been also suggested to be potentially useful in identifying SLRs (McKinney et al., 2020). However, signals of LD have remained under-exploited in population genomic studies. Here we present a method, SLRfinder, which incorporates LD signals to identify candidate SLRs and the sex chromosomes in which they are located. The results show that SLRfinder successfully identified known SLRs as significant candidates when analyzing the published population data of nine-spined sticklebacks and the chum salmon dataset mapped to the male reference genome. In addition, using LD clustering, the SLRfinder-identified SLRs were narrower than those identified using GWAS (Rondeau et al., 2023) or sliding windows (Yi et al., 2024), which indicates that SLRfinder can be beneficial by further narrowing down the highly linked SLR even when the pair of sex chromosomes is already known. Interestingly, the SLRs of nine-spined sticklebacks and chum salmon have been indicated to involve genomic inversions (McKinney et al., 2020; Natri et al., 2019; Yi et al., 2024) which might have strengthened the signals of LD and heterozygosity detected in SLRfinder. Studies have proposed that structural variation, such as inversions and chromosomal fusions, can facilitate recombination suppression in the newly formed SLRs and thus may play important roles in early sex chromosome evolution (Kitano et al., 2009; McKinney et al., 2020; Natri et al., 2019; Yi et al., 2024). However, it remains unclear how often inversions (and other structural variants) might be associated with labile SLRs in natural populations. We propose that SLRfinder might be helpful to answer this question as it is likely most sensitive to the SLRs having structural variants and can be easily applied to genomic data of non-model populations.

We compared SLRfinder, which is based on heterozygosity and mainly developed for labile sex chromosomes, to the previously developed method SATC that is based on the depth of coverage (Nursyifa et al., 2022). As expected, SLRfinder outperformed SATC in analysing labile sex chromosomes that tend to have similar depths between sexes, such as in the WL sticklebacks and chum salmon. In addition, SATC assumes no Y/W-linked scaffolds in the reference genome, which is usually true in the taxa having conserved sex chromosomes because the highly degenerated Y/W chromosomes are difficult to assemble and often excluded when the reference genome comes from the heterogametic sex. However, reference genomes of the taxa having labile sex chromosomes are more likely a mosaic combination of scaffolds from both sex chromosomes if the sequences were from a heterogametic individual (e.g., the version 7 reference of the nine-spined stickleback; Kivikoski et al., 2021), making SATC less applicable and even misleading (such as in the case of EL sticklebacks). In addition, SATC was designed for data mapped to scaffold-level reference genomes (Nursyifa et al., 2022) and could not break down long assembled chromosomes, which may prevent the identification of narrow SLRs of labile sex chromosomes when using chromosome-level reference genomes. On the other hand, SATC worked better than SLRfinder on conserved sex chromosomes having clear inter-sex differences in the depth of coverage. Accordingly, our study suggests that SLRfinder and SATC are complementary methods that specialize on different types of sex chromosomes and datasets (**Table 2**). Therefore, we recommend testing both methods (and potentially other methods as well) when trying to identify SLRs in new populations or species to get complementary results.

**Table 2.** Comparison between SLRfinder and SATC.

|  | **SLRfinder** | **SATC** |
| --- | --- | --- |
| **Recommended type of sex chromosomes** | Labile | Conserved |
| **Considered signals** | Linkage disequilibrium, number of SNPs in the LD cluster, heterozygosity, genetic differentiation | Depth of coverage |
| **Sequencing data** | Whole-genome resequencing (preferred) or restriction site-associated DNA sequencing | Whole-genome resequencing |
| **Reference genome** | Best if chromosome-level; homogametic or heterogametic sex | Best if scaffold-level; homogametic sex (no Y/W-linked scaffolds) |
| **Phenotypic sex** | Not required but can be incorporated to provide extra supports | Not required and not used. |
| **Computational burden** | Medium: memory and speed depend on data size | Small |

Both SATC and SLRfinder do not require phenotypic sexes which can be difficult to obtain from non-invasive sampling or difficult to identify without clear phenotypic sexual dimorphism. Instead, SATC and SLRfinder can be used for genetic sex identification in addition to identifying candidate SLRs and sex chromosomes. Furthermore, SLRfinder also has several advantages as illustrated in our analyses. First, SLRfinder does not require known sex determination systems or a specific reference genome from the heterogametic or homogametic sex. In our study we only tested taxa having the XX/XY system but the same should apply to ZZ/ZW systems. It is worth noting that SLRfinder may work better using the chromosome-level than the scaffold-level reference genome because the former generates more and larger LD clusters. Second, SLRfinder does not require a specific sequencing method (e.g., WGS or RADseq) and can be easily applied to any SNP genotypes in the VCF format. The highly flexible R scripts allow manual parameter settings (e.g., *min_LD*, expected sex ratio, rank parameters) and can be easily extended to include additional ranking or filtering parameters (e.g., *F_ST_* between sexes). Third, SLRfinder is a conservative method. Our test found very few false positives which could be identified by the separation between phenotypic sexes and the usually higher *p*-values than the top-ranked true SLRs. On the other hand, SLRfinder did not have enough power to detect significant SLRs in several cases but the false negatives can be identified by filtering the misplaced sexes. In addition, false negatives tend to be the top-ranked clusters with the lowest non-significant *p*-values and the largest numbers of SNPs.

Like all the other methods, SLRfinder also has its limitations which were explored using the test datasets. First, SLRfinder may have limited power when sample sizes are small, especially with a limited number of populations. For example, SLRfinder successfully identified the true SLR using as few as 24 individuals from eight WL populations of nine-spined sticklebacks, but not when using as many as 79 individuals from 4 WL populations (Table S2). This is likely because more diverse populations generate more and larger LD clusters, which increases the power of SLRfinder. Second, SLRfinder requires sampling both sexes in relatively equal proportions. Although slightly skewed sex ratios (max 1:3 or 3:1) could work in most cases and can be accounted for in ꭓ^2^ tests, SLRfinder appears not to work when sex ratios are highly skewed (e.g., 1:10 or 10:1). However, although not tested here, these limitations from the sample size and sex ratio likely also apply to most of the other methods for SLR identification with few exceptions (e.g., FindZX may work on a single individual; Sigeman et al., 2022).

Unsurprisingly, SLRfinder only works when the expected signals (differential heterozygosity and genetic differentiation between sexes in SLRs) are present in the data. However, these signals may not be clear in every dataset. When sample sizes are small and include low signal-to-noise ratios, these expected signals can occur by chance rather than driven by linkage to sex. In addition, some biological systems may exhibit complicated signals in their SLRs. For example, guppies showed no difference in male and female heterozygosity and stronger population structure than inter-sex divergence in the previously identified candidate SLRs (Fig. 3CD). The high heterozygosity in both sexes and strong population signal might be explained by the maintenance of many different Y haplotypes among these populations via balancing selection (Fraser et al., 2020). Similarly, a previously developed coverage-based method, RADSex, was applied to 15 teleost fishes having labile sex chromosomes but only six were successfully identified with sex markers (Feron et al., 2021). Taken together, these results show that no single method is universally applicable to all taxa having diverse sex chromosome systems.

In summary, SLRfinder provides a novel approach for the identification of labile sex chromosomes in non-model populations using LD and heterozygosity. Given the lack of a universal method for identifying SLRs across diverse sex chromosome systems, SLRfinder complements the previously developed methods (e.g., SATC) by serving the same purpose in different contexts. SLRfinder seems to work best when applied to a large number of divergent populations and when sex ratios are relatively equal. It should be noted that the identified regions are candidate SLRs and putative sex chromosomes which need to be further validated with additional data and other approaches. In addition, SLRfinder is sensitive to inversions in SLRs (e.g., the LG12 and LG3 SLRs in sticklebacks) and can detect autosomal regions that may have become sex-linked (e.g., the LG3 region in chum salmon), which can be interesting in the contexts of sexual selection and antagonism.

## Acknowledgment and Funding

We thank Dr. Bonnie Fraser for the help to get access to the raw data of guppies. This study was supported by National Natural Science Foundation of China/Research Grants Council (RGC) Joint Research Scheme 2021/2022 (‘N_HKU763/21’ to JM). We are grateful for the IT Centre for Scientific Computing (CSC), Finland, for access to computing resources.

## Data availability

SLRfinder scripts are publicly available on GitHub (https://github.com/xuelingyi/SLRfinder) with a step-by-step tutorial. The sample information of our tested datasets are also provided on GitHub.

## Author Contribution

P.K. and X.Y. conceptualized the study. P.K. designed the method and wrote the raw scripts. X.Y. polished the method and analyzed empirical datasets. J.M. supervised the study and provided resources. X.Y. and P.K. drafted the manuscript. All authors edited the manuscript.

## References

Blaser, O., Neuenschwander, S., & Perrin, N. (2014). Sex-chromosome turnovers: the hot-potato model. American Naturalist, 183(1), 140–146. 10.1086/674026

Csardi, G., Nepusz, T., & others. (2006). The igraph software package for complex network research. *InterJournal*, Complex Systems, 1695(5), 1–9.

Danecek, P., Auton, A., Abecasis, G., Albers, C. A., Banks, E., DePristo, M. A., Handsaker, R. E., Lunter, G., Marth, G. T., Sherry, S. T., McVean, G., Durbin, R., & 1000 Genomes Project Analysis Group. (2011). The variant call format and VCFtools. Bioinformatics, 27(15), 2156–2158. 10.1093/bioinformatics/btr330

Danecek, P., Bonfield, J. K., Liddle, J., Marshall, J., Ohan, V., Pollard, M. O., Whitwham, A., Keane, T., McCarthy, S. A., Davies, R. M., & Li, H. (2021). Twelve years of SAMtools and BCFtools. GigaScience, 10(2), giab008. 10.1093/gigascience/giab008

Depristo, M. A., Banks, E., Poplin, R., Garimella, K. V., Maguire, J. R., Hartl, C., Philippakis, A. A., Del Angel, G., Rivas, M. A., Hanna, M., McKenna, A., Fennell, T. J., Kernytsky, A. M., Sivachenko, A. Y., Cibulskis, K., Gabriel, S. B., Altshuler, D., & Daly, M. J. (2011). A framework for variation discovery and genotyping using next-generation DNA sequencing data. Nature Genetics 2011 43:5, 43(5), 491–498. 10.1038/ng.806

Devlin, B., & Roeder, K. (1999). Genomic control for association studies. Biometrics, 55(4), 997–1004. 10.1111/j.0006-341X.1999.00997.x

Devlin, B., Roeder, K., & Wasserman, L. (2001). Genomic control, a new approach to genetic-based association studies. Theoretical Population Biology, 60(3), 155–166. 10.1006/tpbi.2001.1542

Dixon, G., Kitano, J., & Kirkpatrick, M. (2019). The origin of a new sex chromosome by introgression between two stickleback fishes. Molecular Biology and Evolution, 36(1), 28–38. 10.1093/MOLBEV/MSY181

Dufresnes, C., Borzee, A., Horn, A., Stock, M., Ostini, M., Sermier, R., Wassef, J., Litvinchuck, S. N., Kosch, T. A., Waldman, B., Jang, Y., Brelsford, A., & Perrin, N. (2015). Sex-chromosome homomorphy in Palearctic tree frogs results from both turnovers and X–Y recombination. Molecular Biology and Evolution, 32(9), 2328– 2337. 10.1093/MOLBEV/MSV113

Faria, R., Chaube, P., Morales, H. E., Larsson, T., Lemmon, A. R., Lemmon, E. M., Rafajlović, M., Panova, M., Ravinet, M., Johannesson, K., Westram, A. M., & Butlin, R. K. (2019). Multiple chromosomal rearrangements in a hybrid zone between Littorina saxatilis ecotypes. Molecular Ecology, 28(6), 1375–1393. 10.1111/mec.14972

Fang, B., Kemppainen, P., Momigliano, P., Feng, X., & Merilä, J. (2020). On the causes of geographically heterogeneous parallel evolution in sticklebacks. Nature Ecology & Evolution, 4(8), 1105–1115. 10.1038/s41559-020-1222-6

Fang, B., Kemppainen, P., Momigliano, P., & Merilä, J. (2021). Population structure limits parallel evolution in sticklebacks. Molecular Biology and Evolution, 38(10), 4205– 4221. 10.1093/MOLBEV/MSAB144

Feng, X., Merilä, J., & Löytynoja, A. (2022). Complex population history affects admixture analyses in nine-spined sticklebacks. Molecular Ecology, 31(20), 5386–5401. 10.1111/MEC.16651

Feron, R., Pan, Q., Wen, M., Imarazene, B., Jouanno, E., Anderson, J., Herpin, A., Journot, L., Parrinello, H., Klopp, C., Kottler, V. A., Roco, A. S., Du, K., Kneitz, S., Adolfi, M., Wilson, C. A., McCluskey, B., Amores, A., Desvignes, T., … Guiguen, Y. (2021). RADSex: a computational workflow to study sex determination using restriction site-associated DNA sequencing data. Molecular Ecology Resources, 21(5), 1715–1731. 10.1111/1755-0998.13360

Fraser, B. A., Whiting, J. R., Paris, J. R., Weadick, C. J., Parsons, P. J., Charlesworth, D., Bergero, R., Bemm, F., Hoffmann, M., Kottler, V. A., Liu, C., Dreyer, C., & Weigel, D. (2020). Improved reference genome uncovers novel sex-linked regions in the guppy (*Poecilia reticulata*). Genome Biology and Evolution, 12(10), 1789–1805. 10.1093/GBE/EVAA187

Furman, B. L. S., Metzger, D. C. H., Darolti, I., Wright, A. E., Sandkam, B. A., Almeida, P., Shu, J. J., Mank, J. E., & Fraser, B. (2020). Sex chromosome evolution: so many exceptions to the rules. Genome Biology and Evolution, 12(6), 750–763. 10.1093/gbe/evaa081

Guzmán, N. V., Kemppainen, P., Monti, D., Castillo, E. R. D., Rodriguero, M. S., Sánchez-Restrepo, A. F., Cigliano, M. M., & Confalonieri, V. A. (2022). Stable inversion clines in a grasshopper species group despite complex geographical history. Molecular Ecology, 31(4), 1196–1215. 10.1111/mec.16305

Jeffries, D. L., Lavanchy, G., Sermier, R., Sredl, M. J., Miura, I., Borzée, A., Barrow, L. N., Canestrelli, D., Crochet, P. A., Dufresnes, C., Fu, J., Ma, W. J., Garcia, C. M., Ghali, K., Nicieza, A. G., O’Donnell, R. P., Rodrigues, N., Romano, A., Martínez-Solano, Í., … Perrin, N. (2018). A rapid rate of sex-chromosome turnover and non-random transitions in true frogs. Nature Communications, 9(1), 1–11. 10.1038/s41467-018-06517-2

Jeffries, D. L., Mee, J. A., & Peichel, C. L. (2022). Identification of a candidate sex determination gene in *Culaea inconstans* suggests convergent recruitment of an *Amh* duplicate in two lineages of stickleback. Journal of Evolutionary Biology, 35(12), 1683–1695. 10.1111/JEB.14034

Kemppainen, P., Knight, C. G., Sarma, D. K., Hlaing, T., Prakash, A., Maung Maung, Y. N., Somboon, P., Mahanta, J., & Walton, C. (2015). Linkage disequilibrium network analysis (LDna) gives a global view of chromosomal inversions, local adaptation and geographic structure. Molecular Ecology Resources, 15(5), 1031–1045. 10.1111/1755-0998.12369

Kitano, J., Ansai, S., Fujimoto, S., Kakioka, R., Sato, M., Mandagi, I. F., Sumarto, B. K. A., & Yamahira, K. (2023). A cryptic sex-linked locus revealed by the elimination of a master sex-determining locus in medaka fish. The American Naturalist, 202(2), 231–240. 10.1086/724840

Kitano, J., Ross, J. A., Mori, S., Kume, M., Jones, F. C., Chan, Y. F., Absher, D. M., Grimwood, J., Schmutz, J., Myers, R. M., Kingsley, D. M., & Peichel, C. L. (2009). A role for a neo-sex chromosome in stickleback speciation. Nature, 461(7267), 1079– 1083. 10.1038/nature08441

Kivikoski, M., Rastas, P., Löytynoja, A., & Merilä, J. (2021). Automated improvement of stickleback reference genome assemblies with Lep-Anchor software. Molecular Ecology Resources, 21(6), 2166–2176. 10.1111/1755-0998.13404

Kü Nstner, A., Hoffmann, M., Fraser, B. A., Kottler, V. A., Sharma, E., Weigel, D., & Dreyer, C. (2016). The genome of the Trinidadian guppy, *Poecilia reticulata*, and variation in the Guanapo population. PLOS ONE. 10.1371/journal.pone.0169087

Li, H. (2011). A statistical framework for SNP calling, mutation discovery, association mapping and population genetical parameter estimation from sequencing data. Bioinformatics, 27(21), 2987–2993. 10.1093/BIOINFORMATICS/BTR509

Li, H. (2013). *Aligning sequence reads, clone sequences and assembly contigs with BWA-MEM* (arXiv:1303.3997). arXiv. 10.48550/arXiv.1303.3997

Ma, J., & Amos, C. I. (2012). Investigation of inversion polymorphisms in the human genome using principal components analysis. PLoS ONE, 7(7), 40224–40224. 10.1371/journal.pone.0040224

McKinney, G., McPhee, M. V., Pascal, C., Seeb, J. E., & Seeb, L. W. (2020). Network analysis of linkage disequilibrium reveals genome architecture in chum salmon. G3 Genes|Genomes|Genetics, 10(5), 1553–1561. 10.1534/G3.119.400972

Natri, H. M., Merilä, J., & Shikano, T. (2019). The evolution of sex determination associated with a chromosomal inversion. Nature Communications, 10(1), 1–13. 10.1038/s41467-018-08014-y

Nursyifa, C., Brüniche-Olsen, A., Garcia-Erill, G., Heller, R., & Albrechtsen, A. (2022). Joint identification of sex and sex-linked scaffolds in non-model organisms using low depth sequencing data. Molecular Ecology Resources, 22(2), 458–467. 10.1111/1755-0998.13491

Palmer, D. H., Rogers, T. F., Dean, R., & Wright, A. E. (2019). How to identify sex chromosomes and their turnover. Molecular Ecology, 28(21), 4709–4724. 10.1111/MEC.15245

Pečnerová, P., Garcia-Erill, G., Liu, X., Nursyifa, C., Waples, R. K., Santander, C. G., Quinn, L., Frandsen, P., Meisner, J., Stæger, F. F., Rasmussen, M. S., Brüniche-Olsen, A., Hviid Friis Jørgensen, C., da Fonseca, R. R., Siegismund, H. R., Albrechtsen, A., Heller, R., Moltke, I., & Hanghøj, K. (2021). High genetic diversity and low differentiation reflect the ecological versatility of the African leopard. Current Biology, 31(9), 1862–1871.e5. 10.1016/J.CUB.2021.01.064

Perrin, N. (2021). Sex-chromosome evolution in frogs: What role for sex-antagonistic genes? Philosophical Transactions of the Royal Society B, 376(1832). 10.1098/RSTB.2020.0094

R Core Team. (2022). R: A language and environment for statistical computing. R Foundation for Statistical Computing. Vienna, Austria. https://www.R-project.org/.

Rochette, N. C., Rivera-Colón, A. G., & Catchen, J. M. (2019). Stacks 2: Analytical methods for paired-end sequencing improve RADseq-based population genomics. Molecular Ecology, 28(21), 4737–4754. 10.1111/MEC.15253

Rondeau, E. B., Christensen, K. A., Johnson, H. A., Sakhrani, D., Biagi, C. A., Wetklo, M., Despins, C. A., Leggatt, R. A., Minkley, D. R., Withler, R. E., Beacham, T. D., Koop, B. F., & Devlin, R. H. (2023). Insights from a chum salmon (*Oncorhynchus keta*) genome assembly regarding whole-genome duplication and nucleotide variation influencing gene function. G3 Genes|Genomes|Genetics, 13(8). 10.1093/G3JOURNAL/JKAD127

Sigeman, H., Sinclair, B., & Hansson, B. (2022). Findzx: an automated pipeline for detecting and visualising sex chromosomes using whole-genome sequencing data. BMC Genomics, 23(1), 1–14. 10.1186/S12864-022-08432-9/FIGURES/5

Tree of Sex Consortium. (2014). Tree of sex: A database of sexual systems [dataset]. Nature Publishing Group. 10.5061/DRYAD.V1908

Van der Auwera, G. A., Carneiro, M. O., Hartl, C., Poplin, R., del Angel, G., Levy-Moonshine, A., Jordan, T., Shakir, K., Roazen, D., Thibault, J., Banks, E., Garimella, K. V., Altshuler, D., Gabriel, S., & DePristo, M. A. (2013). From fastQ data to high-confidence variant calls: the genome analysis toolkit best practices pipeline. Current Protocols in Bioinformatics, 43(1), 11.10.1–11.10.33. 10.1002/0471250953.BI1110S43

Vicoso, B. (2019). Molecular and evolutionary dynamics of animal sex-chromosome turnover. Nature Ecology and Evolution, 3(12), 1632–1641. 10.1038/s41559-019-1050-8

Yi, X., Wang, D., Reid, K., Feng, X., Löytynoja, A., & Merilä, J. (2024). Sex chromosome turnover in hybridizing stickleback lineages (p. 2023.11.06.565909). bioRxiv. 10.1101/2023.11.06.565909

Zheng, X., Levine, D., Shen, J., Gogarten, S. M., Laurie, C., & Weir, B. S. (2012). A high-performance computing toolset for relatedness and principal component analysis of SNP data. Bioinformatics, 28(24), 3326–3328. 10.1093/BIOINFORMATICS/BTS606

