## Supplementary Materials for "SLRfinder: a method to detect candidate sex-linked regions with linkage disequilibrium clustering"

**Table S1. Summary of the SLRfinder analyses in test datasets.**

**Table S2. Summary of the SLRfinder results in power tests using the stickleback_WL dataset.** The right SLR on LG3 is bolded. The five top-ranked candidates were reported when no significance was found.

**Table S3. Summary of the SLRfinder results in power tests using the stickleback_EL dataset.** The right SLR on LG12 is bolded. The five top-ranked candidates were reported when no significance was found.

* Tables are attached as an independent Excel.

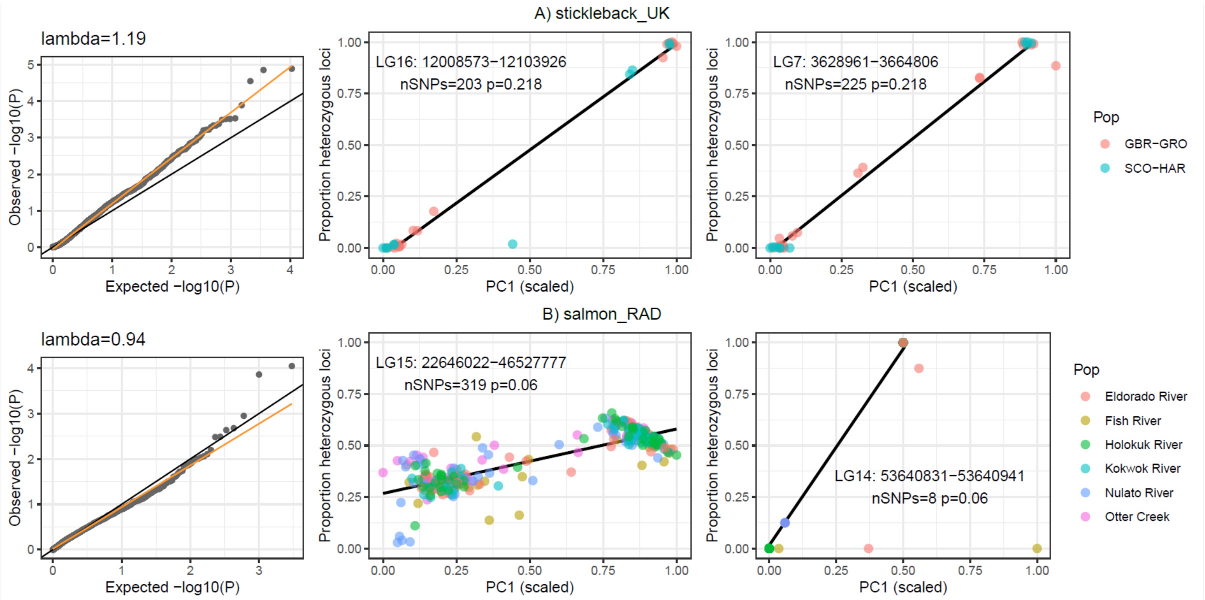


**Figure S1. The SLRfinder-indicated top-ranked candidates in A) the stickleback_UK dataset and B) the salmon_RAD dataset.** The left column shows the QQ plots where dots represent LD clusters colored by significance (no significance found in these datasets). The black line is the expected 1:1 relationship and the orange line is the linear regression fitted to the data points. Regression lines with slopes (i.e., the *lambda* value) >1 indicate *p* value inflation. The right columns are heterozygosity~PC1 plots where dots represent individuals colored by population. **B)** The LG15 region in the salmon_RAD dataset was a false negative. Individuals with scaled PC1 < 0.5 were identified as the homogametic sex (i.e., genetic females in this XX/XY system) and those with PC1 > 0.5 the heterogametic sex (i.e., genetic males in this case). The LG14 region was a true negative with the same adjusted *p* value but only eight SNPs and very few individuals genotyped.

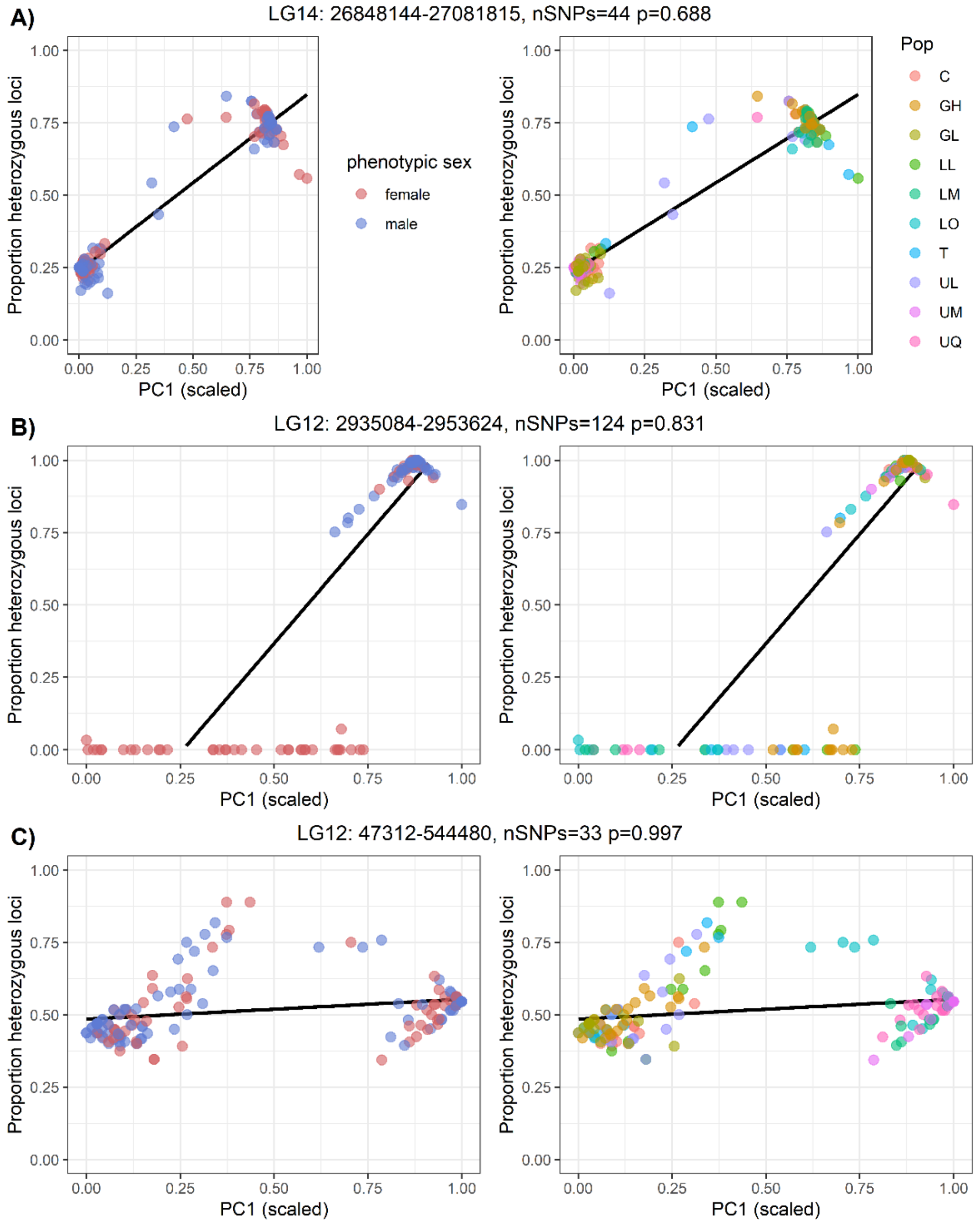


**Figure S2. Heterozygosity~PC1 plots of representative clusters in the guppy_female dataset.** The left column is colored by phenotypic sex and the right column by population. **A)** The SLRfinder-identified top candidate. **B & C)** The two LD clusters identified on the sex chromosomes LG12. Sex was not separated in either of the LG12 clusters. The cluster in C) showed stronger population structure than sex divergence on PC1.

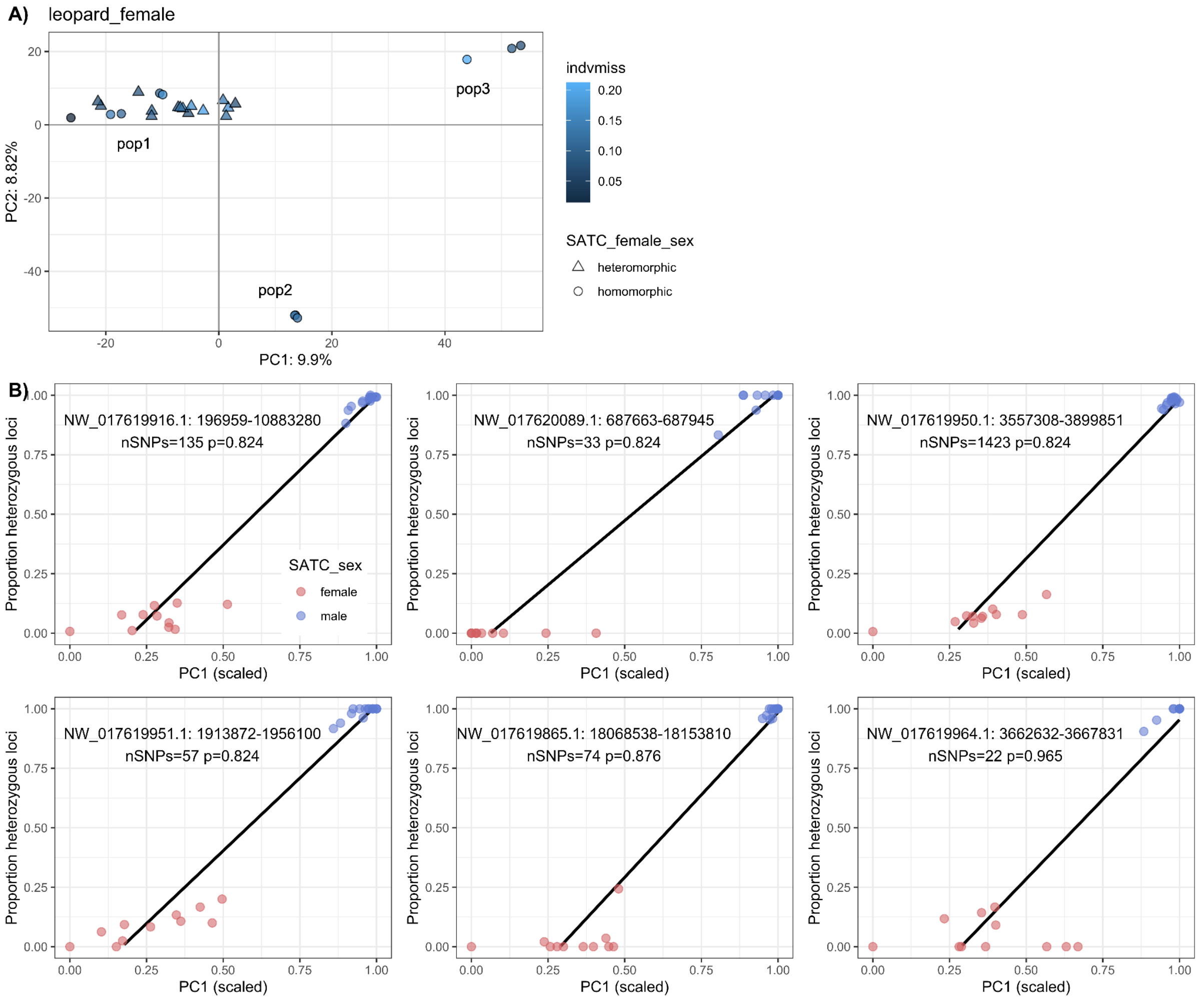


**Figure S3. Analyses of the leopard dataset. A)** The three genetic populations based on PCA. Each dot represents one individual and colors represent individual missing data. The three indicated genetic populations are labeled in text. **B)** The heterozygosity~PC1 plots of sex-separated LD clusters. Colors represent the SATC-inferred genetic sex.
